# OTX2 Dosage Sensitivity is Implicated in Hemifacial Microsomia

**DOI:** 10.1101/001099

**Authors:** Dina Zielinski, Barak Markus, Mona Sheikh, Melissa Gymrek, Clement Chu, Marta Zaks, Balaji Srinivasan, Jodi D. Hoffman, Dror Aizenbud, Yaniv Erlich

**Affiliations:** Whitehead Institute for Biomedical Research, 9 Cambridge Center, Cambridge, MA 02142; Harvard-MIT Division of Health Sciences and Technology, MIT, Cambridge, MA 02139.; Program in Medical and Population Genetics, Broad Institute of MIT and Harvard, Cambridge, Massachusetts, USA; Department of Molecular Biology and Diabetes Unit, Massachusetts General Hospital, Boston, Massachusetts 02114, USA; Counsyl, 180 Kimball Way, South San Francisco, CA 94080; Rambam Health Care Campus, 1 Efron St., Haifa 31096, Israel; Division of Genetics, Tufts Medical Center, Boston, MA 02111

**Keywords:** Hemifacial microsomia, Oculoauriculovertebral spectrum, Exome Sequencing, 14q22 duplication, OTX2, medulloblastoma

## Abstract

Hemifacial microsomia (HFM) is the second most common facial anomaly after cleft lip and palate. The phenotype is highly variable and most cases are sporadic. Here, we investigated the disorder in a large pedigree with five affected individuals spanning eight meioses. We performed whole-exome sequencing and a genome-wide survey of segmental variations. Analysis of the exome sequencing results indicated the absence of a pathogenic coding point mutation. Inspection of segmental variations identified a 1.3Mb duplication of chromosome 14q22.3 in all affected individuals that was absent in more than 1000 chromosomes of ethnically matched controls. The duplication was absent in seven additional sporadic HFM cases, which is concordant with the known heterogeneity of the disorder. To find the critical gene in the duplicated region, we analyzed signatures of human craniofacial disease networks, mouse expression data, and predictions of dosage sensitivity. All of these approaches implicated OTX2 as the most likely causal gene. Moreover, OTX2 is a known oncogenic driver in medulloblastoma, a condition that was diagnosed in the proband during the course of our study. Our findings highlight dosage sensitivity of OTX2 in human craniofacial development and suggest a possible shared etiology between a subtype of hemifacial microsomia and medulloblastoma.

## INTRODUCTION

Hemifacial microsomia (HFM; also termed oculoauriculovertebral spectrum or Goldenhar syndrome, OMIM: 164210) is a highly heterogeneous condition with an estimated rate of 1 in 5,600 to 20,000 births [1]. The hallmarks of this disorder are marked facial asymmetry due to maxillary and mandibular hypoplasia and ear malformations such as preauricular skin tags, microtia, anotia, and conductive hearing loss. Some cases also present epibulbar dermoids and coloboma of the upper eyelid, cleft lip and palate, as well as cardiac, renal, and vertebral defects. To a lesser extent, the disorder also involves neurological anomalies and developmental delays or mental retardation [1–3].

The characteristic facial anomalies of HFM cases are attributed to disruptions in the first and second pharyngeal arches during days 30-45 of gestation in humans [1]. These arches contribute to the development of muscles of mastication, the maxilla, the mandible, middle ear bones, muscles of facial expression, and the stapedial artery. Animal models suggest embryonic hemorrhage or a deficiency in neural crest cell migration as the pathogenesis, disrupting normal development of the pharyngeal arch derived structures [4].

The HFM spectrum reflects a complex pathogenesis that presumably includes both extrinsic and genetic risk factors [2]. Several epidemiological surveys suggest a role for environmental factors that affect the vascular system, including use of vasoactive agents, hypoxia, exposure to teratogens, and gestational diabetes [5]. While most HFM cases are sporadic, approximately 2-10% of cases are familial and present in more than one generation, supporting the contribution of genetic risk factors [6,7]. Careful examination of seemingly unaffected relatives of a large number of probands revealed familial aggregation of mild craniofacial malformations and preauricular skin tags [8]. These mild features are relatively rare in the general population but do not meet the clinical criteria for HFM, leading to a decreased perception of family history. Segregation analysis of 74 families strongly favored an autosomal dominant mode of inheritance with incomplete penetrance over recessive or polygenic transmission [9]. These results suggest that genetics plays a broad etiological role in the manifestation of the disorder.

Genetic investigations of HFM cases have not yet clearly defined the critical genes involved in this disorder. Several studies have reported facial asymmetry and mandibular hypoplasia in cases with gross chromosomal aberrations and trisomies [10–15]. However, these patients exhibit multi-organ pathologies that are atypical of most HFM cases, suggesting that they represent distinct types of syndromes. Genome-wide linkage analysis of 3 HFM pedigrees revealed potential linkage to 14q32, 11q12–13 [16], and 15q26.2-q26.3 [17]. Candidate gene sequencing in these studies failed to find a pathogenic variation. Rooryck et al. [18] performed array CGH on a cohort of 86 HFM patients, most without family history of the disorder. They found 12 copy number variants (CNVs) ranging from 2.7kb to 2.3Mb (median: 153Kb). However, none of these CNVs were recurrent and 9 out of the 10 autosomal CNVs were also present in unaffected individuals. The authors concluded that it is difficult to interpret to what extent these CNVs contribute to the disorder. To date, the field has yet to identify a strong etiological gene that is responsible for the pathogenesis of the disorder.

Here, we conducted a systematic analysis to identify an etiological variant in HFM. To increase the power of the investigation, we focused on a large family with multiple affected individuals. To the best of our knowledge, this family is the largest HFM kinship to date that is described in the literature. We considered both exonic mutations and copy number variations to further increase the probability of identifying the etiological locus while excluding bystander variations [19]. This process revealed a segmental duplication of 8 genes that segregates with the disorder. An unbiased HFM disease network analysis and expression profiling implicate OTX2 as the pathogenic gene in the CNV.

## RESULTS

### Clinical presentation

We identified a five generation Ashkenazi kinship that displays variable HFM anomalies in five individuals separated by a total of eight meiosis events (Figure 1, Table 1). In all cases, the family denied consanguinity and the disorder appears to follow an autosomal dominant segregation pattern with incomplete penetrance and variable expressivity.

**Figure 1:**
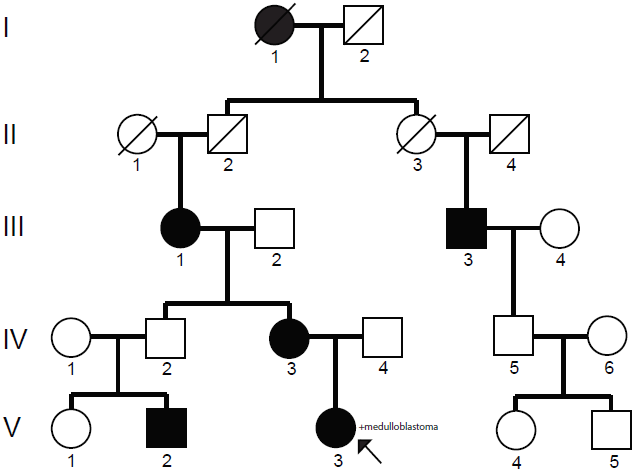
The five-generation pedigree. The family consists of five affected individuals spanning eight meioses. The proband (V.3) is indicated by an arrow.

**Table 1:** Clinical features of family members displaying HFM anomalies

| Clinical feature | III.1 | III.3 | IV.3 | V.2 | V.3 |
| --- | --- | --- | --- | --- | --- |
| Facial cleft | + | + | + | + | + |
| Facial asymmetry | + | + | + | + | + |
| Anotia/microtia | - | - | - | - | + |
| Preauricular tags | + | - | + | + | + |
| Mandibular, maxillary hypoplasia | + | + | + | + | + |
| Retrognathia | - | + | - | + | + |
| Epibulbar dermoids | - | - | - | - | - |
| Cardiac anomalies | - | - | - | - | - |
| Renal anomalies | - | - | - | - | - |
| Vertebral anomalies | - | - | - | - | - |
| Medulloblastoma | - | - | - | - | + |

The proband, subject V.3, was presented to the Craniofacial Department of the Rambam Medical Center in Israel at the age of three. She was born after normal pregnancy (42 weeks) and caesarian delivery. Clinical examination found right mandibular hypoplasia and facial asymmetry, cleft #7 according to Tessier’s craniofacial classification system, preauricular skin tags, and grade II microtia, all on the right side. Deafness in the right ear was diagnosed at the age of 2 months. She is of normal intelligence and no other abnormalities were noted at the time (Table 1). The proband underwent a combined surgical orthodontic manipulation using the distraction osteogenesis technique to elongate the right mandibular ramus. During the course of this study, at age seven, she was diagnosed with a medullosblastoma in the fourth ventricle. The tumor was completely resected, after which the child received craniospinal radiotherapy and chemotherapy [see a case study on her cancer treatment here: [20]].

The proband’s mother (IV.3), grandmother (III.1) and cousin (V.2) were also examined at the Craniofacial Department of the Rambam Medical Center. All individuals exhibited milder facial asymmetry with unilateral clefts and preauricular skin tags without ear involvement. Examination of the proband’s uncle (IV.2) did not reveal any facial anomalies, indicating incomplete penetrance of the disorder.

The proband’s first cousin twice removed (III.3) was identified at a later stage of the study. He presented mild facial asymmetry on his left side without auricle involvement and reported that his grandmother (I.1) displayed similar features.

### Analysis of exonic variants shows evidence of no causal mutation

We performed whole exome sequencing of individuals III.1, V.2, and V.3. The average autosomal coverage of the targeted regions in the three samples was 95x-105x reads per base pair. More than 96% of each exome was covered by at least one read (Supplementary Figure 1). Exome sequencing revealed 22,252, 22,746, and 23,175 exonic variants in III.1, V.2, and V.3 respectively. We observed transition/transversion ratios of 2.89-3.00 and homozygous to heterozygous mutation ratios of 0.56-0.58. In parallel, we also conducted genome-wide genotyping of these three samples using the Affymetrix SNP Array 6.0. Comparing shared variations between the two platforms showed concordance rates of more than 98% for non-reference loci (Supplementary Table 1). All of these technical indicators are consistent with the results of previous studies [21–23], supporting the quality of the exome sequencing data.

We passed the exonic variations through a series of filters to find mutations that fit the rare familial pathology (Table 2). First, we excluded synonymous variants. Second, we excluded variations that appear at a frequency greater than 0.1% in large-scale sequencing projects such as the Exome Sequencing Project, 1000 Genomes, and ClinSeq, as documented in dbSNP. In addition, we excluded variations that also appeared at least twice in the exome sequencing data of 21 healthy Ashkenazi Jews (provided by Noam Shomron, Tel Aviv University). In the **Supplementary Note**, we show that these frequency cutoffs are very conservative. Third, we focused only on variants that reside in regions that are identical by descent (IBD) in all individuals. Variants that reside in these haplotypes where transmitted from III.1 to V.2 and V.3. Shared variants outside these regions are from ancient coalescent events and reflect inheritance patterns that do not segregate with the phenotype. Using genome-wide genotype data, we identified 33 autosomal segments that are IBD in these three individuals, with a total size of 421.2Mb (14.5% of the autosome). This value is close to the theoretical expectation of a familial relationship of one grandmother and two cousins (1/4×1/2=12.5% on average). After excluding exonic variations that fall outside these segments, the number of plausible candidates was reduced to 84, 90, and 72 variations in III.1, V.2, and V.3. Finally, we retained only variations in the IBD segments that appear in all three individuals (Supplementary Table 3), which resulted in 40 candidates (26 SNPs and 14 indels). Only 4 out of these 40 variations were not documented in dbSNP.

**Table 2:** Exome filtering steps

| Filtering steps | III.1 | V.2 | V.3 |
| --- | --- | --- | --- |
| Exonic variants | 22,252 | 22,746 | 23,175 |
| Non-synonymous | 9,552 | 9,839 | 10,072 |
| Rare variants | 560 | 662 | 665 |
| Variants in IBD segments | 84 | 90 | 72 |
| Shared variants | 40 |  |  |
| Shared with III.3 | 0 |  |  |

At this stage, we were able to recruit individual III.3 to the study. We conducted array-based genome-wide genotyping and used the results to determine shared segments that are IBD in all four individuals: III.1, III.3, V.2, and V.3. This process resulted in 16 segments with a total length of 59Mb (2.0% of the autosome that is shared between all four individuals). Again, this number is close to the theoretical expectation of 1/4×1/4×1/4=1.6%. Excluding variants outside these regions returned *zero* shared candidates of the 32 variants from the previous step. This filtering process showed that there is no single non-synonymous variant of relatively rare frequency in the population that segregates with the disorder.

To further validate our findings, we performed Sanger sequencing of 37 variants that were identified in the exome sequencing results but excluded after the final IBD filtration step. Four of these variants were located in genes with biological activities that could relate to the disorder (DAB2, IQSEC1, KIAA1456, and ADAM28), such as vascularization, angiogenesis, imprinting, and neurogenesis [24–27]. However, Sanger sequencing of all 37 variations, including these four genes, showed that individual III.3 is does not carry the variant, as expected from the IBD analysis (Supplementary Figure 2; Supplementary Table 4). Importantly, these results support the validity of the IBD filtration technique and provide additional evidence supporting the absence of an etiological point mutation in the exome.

### Copy Number Variation Analysis Identifies a Familial Duplication of 14q22.3

Given the absence of point mutations, we turned to copy number analysis using the genotype data from the genome-wide SNP array. Our analysis revealed a 1.3 Mb duplication of 14q22.3 (chr14:57,141,867-58,495,517) in all four individuals that segregated along all 8 meioses (Figure 2a). In general, CNVs of this length are rare and typically deleterious [28]. No other detected CNVs (>10kb) were found to segregate with the disorder. To increase the sensitivity, we repeated the CNV analysis and inspected only CNVs that are shared in individuals III.1, V.2, V.3. We excluded individual III.3 from this analysis because the array genotyping was performed separately and showed greater systematic noise. This process revealed seven CNV segments (>10Kb) in addition to the duplication of 14q22.3. However, all but one where also found in healthy Ashkenazi controls from genome-wide genotyping array data [29]. The one segment that was not present in the Ashkenazi controls was a ∼37 kb duplication of a non-coding region (chr3:187,279,170-187,316,070) that overlapped a known duplication found in healthy Asian controls in the Database of Genomic Variants (DGV: nssv1548729). Moreover, we did not see any evidence of this region in the array data for III.3. Thus, we concluded that the duplication of 14q22.3 is the only likely CNV that segregates with the disorder.

**Figure 2:**
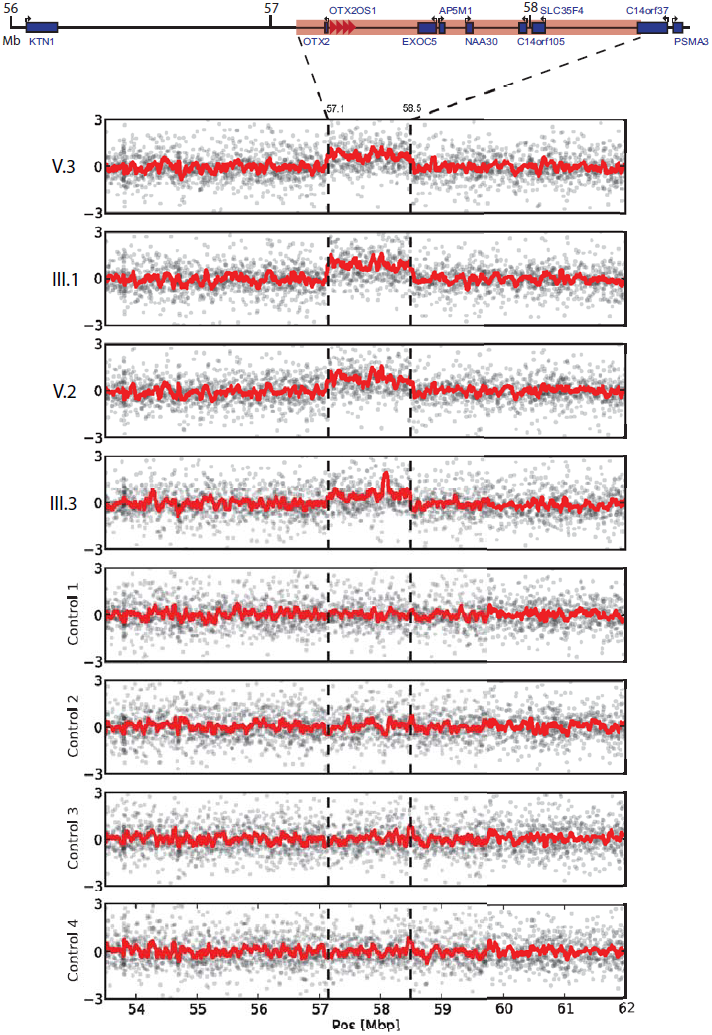
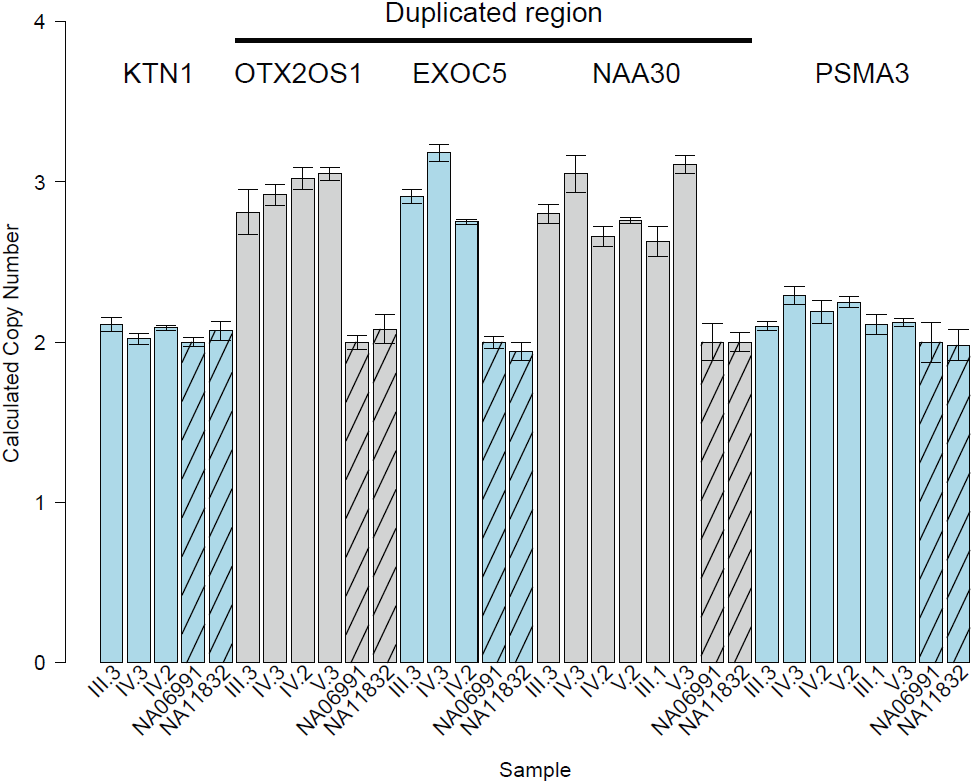
The 14q22 duplicated region. (a) Raw intensity plots of the duplicated region (contained between the dotted lines) in the four affected individuals and 4 Askhenazi controls from (Bray et al. 2010). The signals represent the number of standard deviations of the probes from the mean value. The suspected copy number gain is marked by dotted vertical lines. The red line is a moving average with a window of 20 probes. (b) qPCR results of the affected family and two HapMap controls for genes in the duplcated region (OTX2OS1, EXCO5, and NAA30) and two flanking genes (KTN1 and PSMA3) are consistent with the array results.

In order to confirm the expected rarity of this duplication, we evaluated its frequency in the general Ashkenazi population. Analysis of the genome-wide genotyping array data from 942 healthy Ashkenazi chromosomes [29] returned two copies for this region. In addition, no duplications were found in this region in CNV analysis of deep whole genome sequencing data from 284 chromosomes of Ashkenazi controls sequenced by Complete Genomics that are part of The Ashkenazi Genome Consortium (TAGC) and 1842 chromosomes from phase I of the 1000 Genomes Project [30]. These population-specific results support a familial variant that segregates with the disorder.

To validate our results, we performed qPCR analysis of the duplicated region using Taqman assays (Figure 2b). Three probes targeting genes in the duplication (OTX2-OS1, EXOC5, and NAA30) were confirmed as CN=3 (copy number) in individuals IV.2, IV.3, and III.3. We also observed duplication of OTX2-OS1 and NAA30 in V.3 and of NAA30 in III.1, confirming segregation of this CNV along all informative meioses of the family. Assays targeting OTX2-OS1, EXOC5, and NAA30 returned CN=2 in all HapMap controls and OTX2-OS1 and NAA30 were both CN=2 in 45 Ashkenazi control samples. To validate the boundaries of the CNV, we also targeted KTN1 and PSMA3, upstream and downstream of the predicted CNV. Both probes returned CN=2 in affected family members and HapMap controls (Figure 2b).

In order to evaluate the presence of the duplication in additional HFM cases in Israel, the Craniofacial Department of Rambam Medical Center collected DNA from 7 families that consisted of one affected offspring and unaffected parents. Interrogation of 2 genes in the duplicated region (NAA30 and OTX2-OS1) by qPCR did not reveal any copy number changes in the seven additional HFM cases (Supplementary Figure 3). These findings suggest a distinct genetic etiology of the disorder in our family and are consistent with previous studies that described genetic heterogeneity [18]. However, a literature search revealed that a spectrum of genetic lesions in the 14q22 region have been associated with various facial anomalies. Ou et al. [31] reported a complex event of a duplication of 11.8Mb that fully encompasses our 14q22 region and translocation to 13q21. Interestingly, the proband suffered a range of clinical signs resembling HFM including facial asymmetry, mandibular hypoplasia, and ear defects in addition to developmental delay, lacrimal duct stenosis and renal anomalies. Northup et al. [32] reported a large pericentric inversion inv(14)(p11.2q22.3) in a proband with HFM signs, inherited from his phenotypically normal mother. Ballesta-Martinez et al. [33] recently published a short clinical report of a 14q22 duplication in a Spanish family with variable phenotypes resembling HFM. All of these add additional support to our findings.

### Candidate Gene Prioritization in the Duplicated Segment

We sought to predict the etiological gene that contributes most to the phenotype in an unbiased manner among the eight genes (OTX2, OTX2-OS1, EXOC5, AP5M1, NAA30, C14orf105, SLC35F4, and C14orf37 [partial]) that reside in the duplicated region.

First, we prioritized the genes in the duplicated region based on the similarity of their molecular signatures with known etiological genes of other facial malformations. We and others have successfully identified etiological genes using this guilt-by-association approach in previous studies of rare human disorders [34–36]. The basis of this technique is that similar phenotypes are caused by genes that reside in close biological modules, such as the same pathway, co-expression cluster, and shared regulatory control (Goh et al 2007). To identify a set of disorders similar to HFM in an unbiased manner, we used MimMiner, which ranks clinical conditions in OMIM based on phenotypic resemblance [37]. The top three phenotypes with similar features to HFM were CHARGE syndrome (OMIM: 214800), VACTERL association (OMIM: 314390), and Townes-Brocks syndrome (OMIM: 107480). In fact, HFM and TBS are both characterized by first and second arch defects, including ear, jaw, and kidney malformations [38]. Interestingly, a previous study also cited the commonalities between HFM, CHARGE, and VACTERL [39], adding additional support to the MimMiner prediction. We then compared the biological signatures of all coding genes in the duplicated region to CHD7, ZIC3, and SALL1 the corresponding genes of the three syndromes. To increase the robustness of our analysis, we tested these similarities using two gene prioritization tools: Endeavour [40] and ToppGene [41]. These algorithms utilize different biological datasets and employ distinct prioritization procedures. These two algorithms independently ranked OTX2 as the gene with the closest molecular signature to other facial anomalies (Figure 3a).

**Figure 3:**
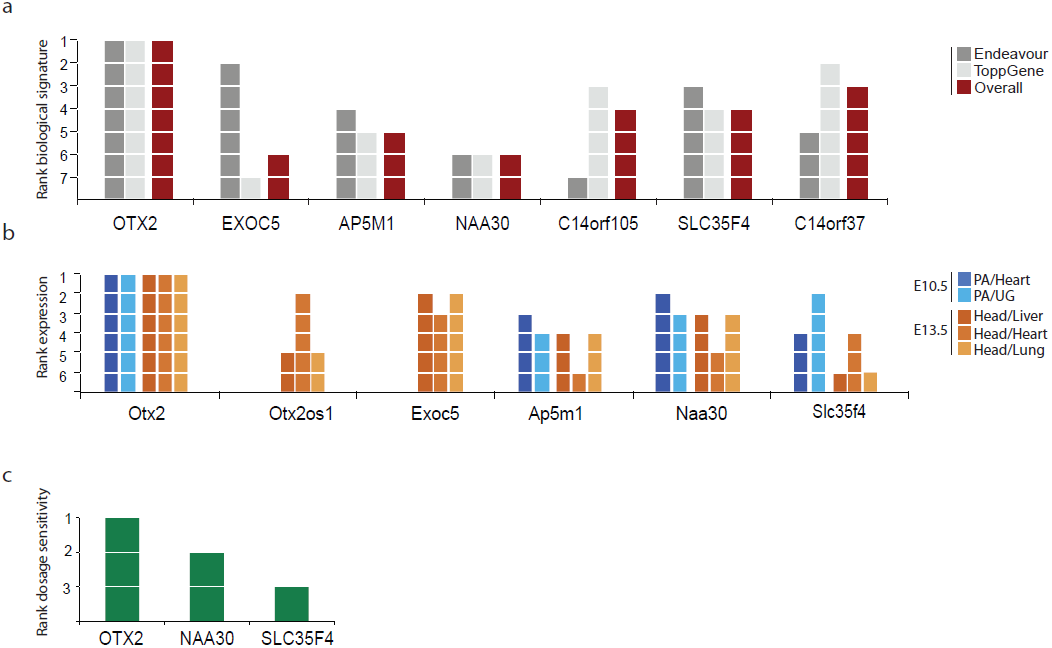
Prioritization of genes in 14q22. (a) Ranking of expression levels in pharyngeal arches (PA) compared to heart and urogenital epithelium (van Driel et al. 2006) in E10.5 and expression in the head compared to liver, heart, and lung in E13.5 for genes in the duplicated region. Comparative expression ranked OTX2 highest in the affected tissues in all conditions (b) Ranking similarity of the molecular signatures of the genes in the duplicated region to causal genes in CHARGE, VACTERL, and Townes-Brocks using Endeavour and ToppGene. The average rank of both toolsis indicated in red. c) Ranking of dosage sensitivity predictions for 3 of the duplicated genes (Huang et al. 2010).

Disease genes tend to be more highly expressed in affected tissues than in those that are unaffected [42,43]. In order to further support the pathogenicity of the duplication, we used publicly available expression array profiles of mouse embryonic tissue to compare the expression of the duplicated genes in affected versus unaffected tissues. Specifically, we analyzed expression levels in the pharyngeal arches at embryonic day 10.5 and in the entire head at E13.5. These developmental stages approximately overlap with the suggested critical periods for the HFM developmental perturbation in humans [1]. We contrasted these expression levels with the expression profiles of liver, heart, and lung (E10.5) and heart and urogenital epithelium (E13.5) since these tissues are rarely implicated in HFM. At E10.5, the arrays contained data for Otx2, Ap5m1, Naa30, and Slc35f4. At E13.5, the arrays contained data for Otx2, Otx2os1, Exoc5, Ap5m1, Naa30, and Slc35f4. The expression profiles showed that Otx2 tends to be more highly expressed in the affected tissues than other duplicated genes at E10.5 and E13.5 compared to any of the unaffected tissues (Figure 3b).

Finally, we also evaluated the general sensitivity of the genes in the region to duplication. Huang et al. [44] developed a gene-level classifier that compares evolutionary, functional, gene-structure, and interaction patterns between haplosufficient and haploinsufficient genes. Interestingly, they found higher expression and tissue specificity of haploinsufficient genes early in development. Although the classifier predicts the probability of haploinsufficiency, it is also useful for detecting genes with increased dosage sensitivity (M. Hurles, personal communication, August 2013). Three of the duplicated genes were included in their classifier: OTX2 had the highest sensitivity score (0.9) followed by NAA30 (0.474) and SLC35F4 (0.418) (Figure 3c). To summarize, all of our *in silico* analysis techniques suggested that duplicated OTX2 is the most likely pathological gene in our HFM cases.

## DISCUSSION

We conducted a systematic study of familial HFM that implicates OTX2 dosage sensitivity in the disorder. OTX2 encodes a transcription factor that plays a critical role in craniofacial development and anterior brain morphogenesis. Loss-of-function studies in mice showed that null embryos fail to develop the anterior head and die during embryogenesis while *Otx2*^+/−^ mice exhibit a range of severe craniofacial anomalies, including micrognathia, agnathia, anophthalmia, and head narrowing [45]. The severity of the phenotype depends on the genetic background [46], consistent with the wide spectrum of phenotypes associated with loss of function in humans. Temporal loss of one copy of Otx2 during mouse embryogenesis up to E12.5 results in haploinsufficiency that leads to significantly low survival rates and abnormal head development, including reduction or absence of the forebrain, eyes, and jaw [47]. OTX2 hemizygous deletions and non-synonymous point mutations have been reported in patients with severe ocular malformations and hypopituitarism, symptoms that are not seen in our pedigree [48–50].

The OTX2 germline duplication in our case suggests a potential link to the medulloblastoma of the proband. OTX2 is a known oncogenic driver of medulloblastoma [51]. Focal duplications and overexpression of this gene are prevalent in subclasses C and D of medulloblastoma [52]. Analysis of her tumor revealed an additional loss of heterozygosity on chromosome 17q [20] that is exclusively associated with subclasses C and D [52]. The potential biological link between OTX2 duplications in hemifacial microsomia and medulloblastoma raises the possibility of their comorbidity. While confirming this hypothesis will require the analysis of a large number of cases, we suggest clinicians be aware of the possibility of increased risk for medulloblastoma in HFM cases with OTX2 duplications.

Our study adds to the existing literature in multiple ways. First, our study considers the largest HFM pedigree to date, increasing the confidence of our genetic analysis. Second, it is the first HFM study to combine whole exome sequencing analysis with the scanning of copy number variants. This approach increases the likelihood that the duplicated region is indeed the etiological site. Third, we present data from more than 1000 chromosomes of unaffected controls, which strongly diminishes the likelihood that the duplication is a polymorphism that segregates in the population. Fourth, we report an unbiased search using different systems biology approaches to find the most likely pathological gene in the region. These analyses implicated OTX2 as the most likely causal gene. Fifth, our findings suggest a potential shared etiology for HFM and medulloblastoma.

Determining the causative gene for HFM can promote stratification of cases based on the molecular pathology, guide clinical care, offer reproductive alternatives to families that carry an OTX2 duplication, and facilitate definitive diagnosis, which is currently inadequate for HFM. Importantly, implicating OTX2 in this disorder can improve understanding of the basic molecular processes that underlie normal and pathological craniofacial development.

## MATERIALS AND METHODS

### Human Subject Research

This study was approved by the Helsinki Committee at the Rambam Medical Center (Haifa, Israel), the Israeli Ministry of Health, and MIT’s COUHES committee.

### Coordinate System

All alignment and genomic coordinates in this manuscript are reported according to hg19. All coverage values are reported after removing PCR duplicates.

### DNA Collection

All DNA was derived from whole blood using standard procedures.

### Exome Sequencing

Paired-end library preparation and exome enrichment were done following a streamlined protocol written by Blumenstiel et al. [53], using Agilent’s SureSelect All Exon V.2 kit, which covers 98.2% of exons and splice sites, according to the Consensus CDS (CCDS) database [54]. Sequencing was performed at Counsyl (South San Francisco, USA) on a single flow cell on the Illumina HiSeq2000 with 100 bp paired end reads (V.2 and V.3 on 3 lanes and III.1 on 2 lanes).

To increase the accuracy of our analysis, we processed the sequencing data with two distinct pipelines. First, we iteratively aligned the sequence reads with Bowtie [55] and with BWA [56]. Multi-mappers were excluded. Reads that failed to align were repeatedly trimmed by 10bp down to a minimum of 36 bp and were processed in an additional round of alignment. The BAM files of all unique mappers from the different alignment rounds were merged and PCR duplicates were removed using SAMtools [57]. Variant calling of Bowtie-aligned reads was done using VarScan v2.8.8 [58] with mpileup2cns and the following options: --min-coverage 5 --min-freq-for-hom 0.9 --p-value 0.97 --strand-filter 1. After alignment using BWA, variant calling was done using the Genome Analysis Toolkit (GATK) [59], following the recommended workflow and filtering of low quality variant calls. In addition, we used lobSTR 1.0.6 [60] to examine short tandem repeat variations in the exomes of III.1. V.2, and V.3. We filtered for STRs genotyped in all three samples with at least 5x coverage in each, that fell within regions shared by all samples with IBD=1, and falling within annotated Refseq genes. Six loci were called as non-reference in all three samples. For each locus, the non-reference allele was found in at least one healthy control from a panel of more than 30 healthy controls, mainly of European descent.

### Validation by Sanger Sequencing

We used Primer3 [61] to design primers flanking candidate variants (+/−100bp upstream and downstream). We excluded primers that generated more than one *in silico* pcr product on the UCSC Genome Browser [62]. Sanger sequencing was done on an ABI 3730 DNA Analyzer.

### Genome-Wide Human SNP Array 6.0

Genomic DNA was extracted from peripheral blood leukocytes using standard methods. We performed genotyping of subjects III.1, III.3, V.2, V.3 using the Affymetrix SNP 6.0 Array. We analyzed the 4 cases together with 471 unrelated Ashkenazi controls [29] (NCBI GEO GSE23636) using the Affymetrix genotyping console (v 4.1.3) and Birdsuite [63] for genotype calling.

### Investigating exonic variations

Annotation of exonic variations was done using SeattleSeq 137 [23] and minor allele frequencies in dbSNP were taken from BioQ [64]. Filtering of variants was done using BEDTools [65] and custom Perl scripts (available upon request).

### IBD Calculations

We used the Affymetrix genotyping console (v 4.1.3) for genotype calling of our 4 subjects together with 50 randomly selected individuals from the Ashkenazi controls (Bray 2010). Initial data analysis and selection of SNPs were carried out using PLINK [66]. We selected subsets of SNPs with MAF > 0.1 that are in approximate linkage equilibrium. This was carried out using the pairwise correlation method for LD pruning implemented in PLINK. We used the following parameters: window size = 50, step = 5, r＾2 threshold = 0.35. The pruned data contained 123209 SNPs.

We used the pruned data as input to MERLIN [67] for pairwise IBD inference, with genetic map positions of 1Mbp = 1cM. Candidate IBD regions were selected based on pair-wise IBD probabilities. We marked all regions for which IBD probabilities for sharing an allele for all pairs of cases in the data were inferred to be higher than 0.5. We then extended the IBD region to include the tips of the chromosomes for cases when IBD = 1 was detected in the first or the last SNP on the chromosome.

### Taqman CNV Assays

We purchased custom Taqman probes to interrogate the CNV and flanking regions (probe start locations in NCBI build 37: chr14:20811565, chr14:56099993, chr14:57267695, chr14:57270923, chr14:57272149, chr14:57277101, chr14:57328402, chr14:57476529, chr14:57597148, chr14:57700715, chr14:57868427, chr14:58725337, chr17:44203062). Reactions were carried out in 10ul, with 10ng genomic DNA and 10ng reference DNA (RnaseP), in 4 replicates. Copy number was determined using the delta delta Ct method and CopyCaller v2.0 with HapMap samples NA06991 and NA11832 as calibrators. The OTX2 probes that were purchased from ABI failed to work despite repeated attempts. They produced non-Mendelian inheritance patterns for trios and reported deletions of the region in normal healthy controls. We therefore excluded these probes from the analysis.

### Prioritization using Biological Signatures

Endeavour is available at: http://homes.esat.kuleuven.be/∼bioiuser/endeavour/tool/endeavourweb.php and ToppGene is available at: http://toppgene.cchmc.org/prioritization.jsp. In Endevaour, we used the following features: CisRegModule, Expression - SonEtAl, Expression - SuEtAl, Interaction - Bind, Interaction - BioGrid, Interaction - Hprd, Interaction - InNetDb, Interaction - Intact, Interaction - Mint, Interaction - String, Motif, Precalculated - Ouzounis, and Precalculated - Prospectr. In ToppGene, we used the following features: Domain, Pathway, Interaction, Transcription Factor Binding Site, Coexpression, Computational, MicroRNA, Drug, and Disease.

### Expression analysis of genes in the region

Expression profiles were derived from the following experiments in GEO [68]: Pharyngeal arches E10.5: experiment GDS3803 with subjects GSM448013, GSM448014, GSM448015, GSM448016, and GSM448017. Urogenital epithelium E10.5: experiment GDS3173 with subjects GSM257875, GSM257932, and GSM257933. Heart E10.5: experiment GDS627 with subjects GSM25150, GSM25151, GSM25152. Head E13.5: experiment GDS2874 with subjects GSM212558, GSM212560, GSM212562, and GSM212564. Liver E13.5: experiment GDS2693 with subjects GSM177034, GSM177035, and GSM177036. Lung E13.5: experiment GSM290632 with subject GSE11539. All experiments were done using the Affymetrix Mouse Expression Array 430. The pharyngeal arches experiment reported results only from the A array and all the others reported both the A and B arrays. Therefore, in all E10.5 comparisons, we restricted the analysis only to genes that are on the A array.

Based on experimental details in GEO or associated publications, the genetic background of all mice was concluded to be C57BL/6, with the exception of GDS3173 (E10.5 urogenital epithelium), the background of which was not documented.

We downloaded the full soft file of each experiment from GEO, extracted the data from the relevant subjects, and normalized the expression data to range from zero to one for each subject. Experiments with multiple sets were averaged inside the same condition. Then, genes with more than one probe were averaged inside the same condition. Finally, we divided the expression of each gene in the affected tissue (pharyngeal arches and head) by expression in the control tissues (liver, lung, heart, and urogenital epithelium) and ranked the expression levels.

### Dosage sensitivity analysis

Data was taken from Dataset_S1.txt of Huang et al. [44].

## ACKNOWLEDGEMENTS

We gratefully acknowledge the study participants and thank Noam Shomron, Itsik Pe’er, Bob Handsaker, and Sara Selig for providing valuable information about the duplicated region in control samples. We also thank the Whitehead Institute’s Genome Technology Core for assistance in producing the array datasets. YE is an Andria and Paul Heafy Family Fellow and holds a Career Award at the Scientific Interface from the Burroughs Wellcome Fund. This study was also supported by generous gifts from Cathy and Jim Stone and Ron Casty.

## Supplemental Note

Our working hypothesis was that any point mutation that causes HFM will have a minor allele frequency (MAF) of less 0.1% in large sequencing projects. We based our hypothesis on the fact that HFM is estimated to occur at a frequency of 1:5,000-1:20,000 births in the general population. Segregation analysis by Kaye et al. (1992) predicted that the **sum** of minor allele frequencies of all HFM causative genes is 1:3000 (after taking into account penetrance levels). The MAF of a **single** etiological variant is even smaller, since previous linkage analysis identified at least three non-overlapping segments.

Moreover, the affected family is of Ashkenazi heritage. With the limited gene flow between the Ashkenazi population and other European populations, the causal mutation in our family is expected to be at even smaller frequencies in these large sequencing projects due to the low sampling rates of Ashkenazi Jews. To confirm this assumption, we compared the MAFs of more than 50 recessive mutations associated with Ashkenazi genetic disorders to the Exome Sequencing Project where we obtained most of the control chromosomes used in our analysis. These mutations are found at frequencies of 1/25 to 1/70 in the Ashkenazi population, which is much higher than the expected frequency of a causative mutation of HFM. We found that the MAFs of these mutations were diluted by factors of more than 20x to 50x in ESP compared to the Ashkenazi population. Even if the causal mutation is found at a very unlikely rate of 1% in Ashkenazim, we expect it to be <0.05% in ESP. Thus, a 0.1% threshold is highly unlikely to miss the causative mutation.

Similarly, we excluded variants that were seen at least twice in 42 unaffected Ashkenazi chromosomes. The probability to see a mutation with a true MAF of 0.1% in two individuals from this cohort is < 1×10^−3^. Therefore, there is a very small risk of excluding the causative mutation using this MAF cutoff.

**Supplemental Figure 1:**
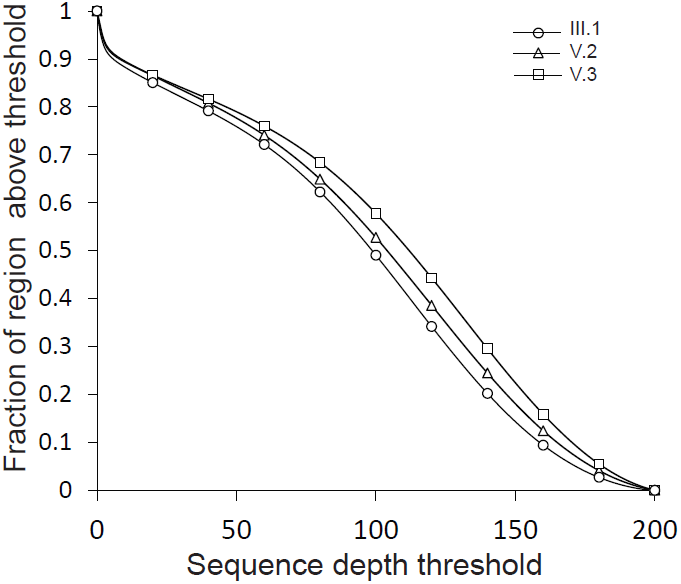
Distribution of exome sequencing cover-ages for the three datasets.

**Supplemental Figure 2:**
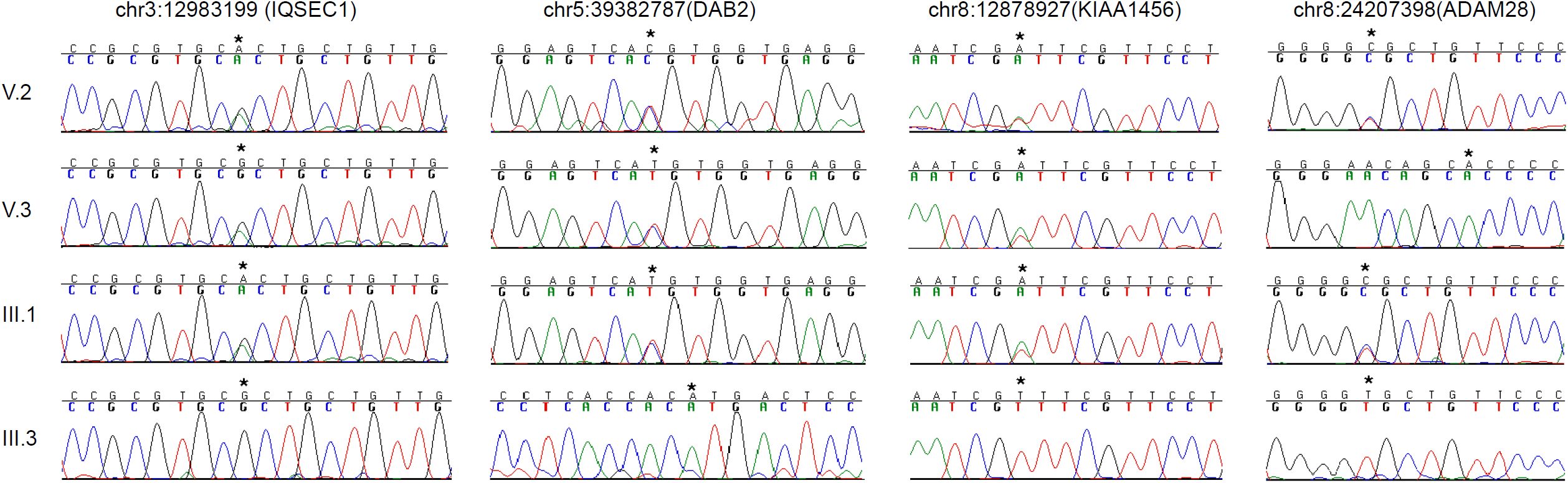
Sanger traces of the four genes with biological activity that could be associated with HFM.

**Supplemental Figure 3b:**
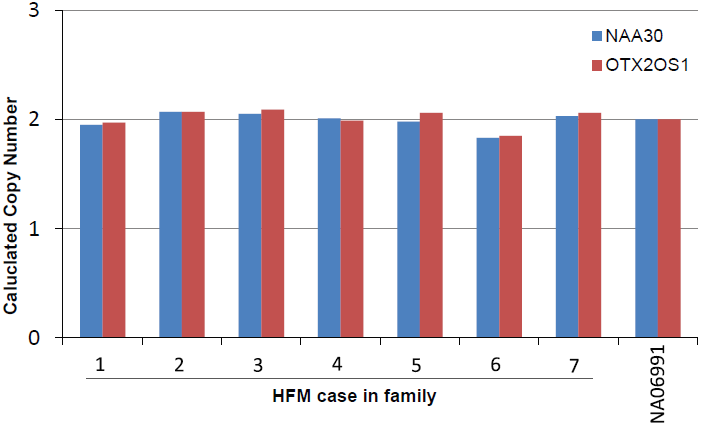
qPCR results of the probands in the seven families. Both tested probes show copy number 2 of the critical region. NA06991 is a HapMap control.

**Supplemental Figure 4:**
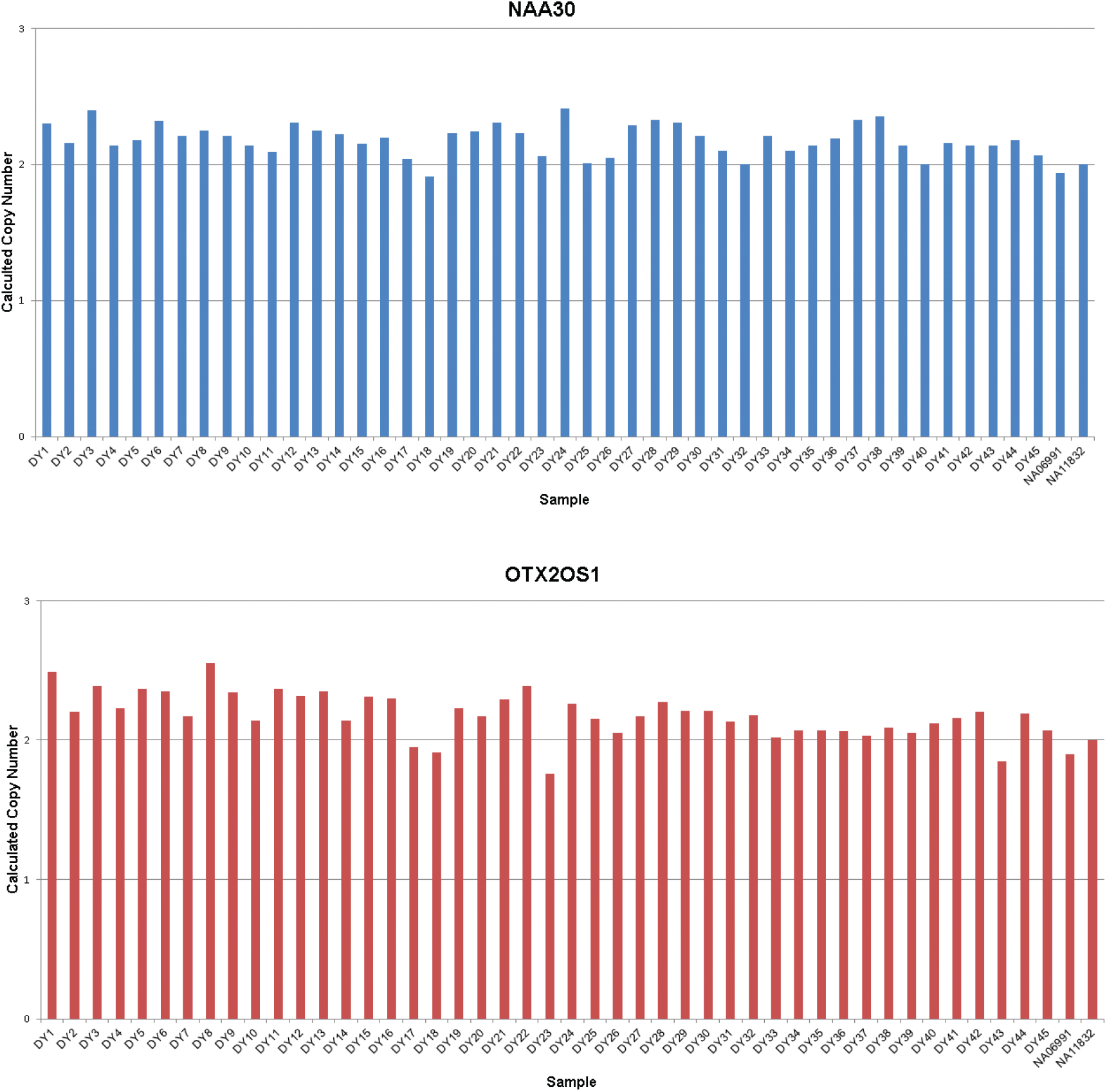

**Supplementary Table 1:**
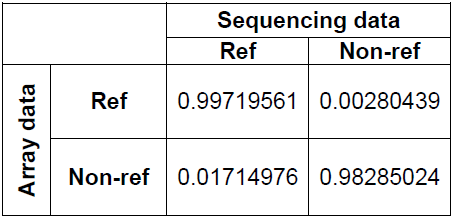
The probabilities to observe genetic variants in the sequencing data conditioned on the array data status and collapsed in all three individuals.

**Supplementary Table 2:** A summary of the quality control indicators from the three exome sequencing datasets.

|  | III.1 | V.2 | V.3 |
| --- | --- | --- | --- |
| # lanes (Illumina) | 2 | 3 | 3 |
| Average coverage | 95.32x | 99.24x | 105.26x |
| Exome covered | 96.00% | 97.30% | 96.90% |
| Exome covered $\geq 5x$ | 90.40% | 91.60% | 91.30% |
| Ts/Tv rate | 3 | 2.89 | 2.93 |
| Homozygous:heterozygous | 0.57 | 0.58 | 0.56 |

**Supplementary Table 3:** Variants shared IBD in individuals III.1, V.2, and V.3.

| chr | pos | observed alleles |
| --- | --- | --- |
| 1 | 1334409 | C/G |
| 1 | 1900106 | insCCT |
| 1 | 229462617 | G/T |
| 2 | 129075877 | G/T |
| 2 | 130832185 | A/A |
| 2 | 130832292 | A/T |
| 3 | 12983199 | A/G |
| 3 | 56650051 | insCTT |
| 8 | 6673377 | -A |
| 8 | 6679498 | G/T |
| 8 | 7308386 | C/C |
| 8 | 7673126 | A/C |
| 8 | 8887542 | delAAC |
| 8 | 10467652 | C/G |
| 8 | 11995570 | G/T |
| 8 | 12878927 | A/T |
| 8 | 64098729 | insG |
| 8 | 86126827 | insAACATT |
| 9 | 894197 | G/T |
| 9 | 21077767 | C/G |
| 10 | 97920099 | insC |
| 10 | 118383463 | insG |
| 10 | 126683123 | A/C |
| 10 | 126683151 | C/T |
| 12 | 7080210 | insG |
| 12 | 7456988 | C/T |
| 12 | 8327883 | C/T |
| 12 | 8374781 | -/ACG |
| 12 | 9994445 | delTGT |
| 12 | 10332200 | A/G |
| 12 | 10573094 | C/G |
| 12 | 10588530 | C/G |
| 12 | 11149585 | A/C |
| 12 | 11244149 | A/G |
| 12 | 11420333 | -/G |
| 12 | 11506669 | G/T |
| 12 | 18435398 | -/CCC |
| 12 | 55523586 | -T |
| 12 | 57433048 | C/T |
| 12 | 75816814 | insACA |
| 15 | 22074657 | C/G |

**Supplementary Table 4:**
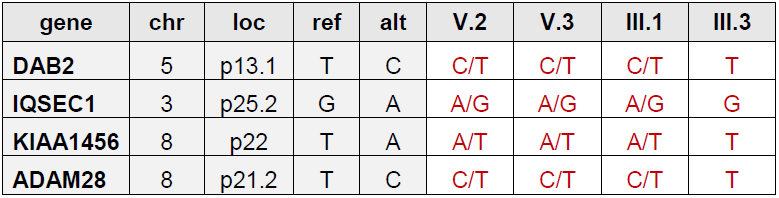
The Sanger sequencing results of the four genes with biological activity that could be attributed to HFM.

## REFERENCES

1. Gorlin RJ CM, Hennekam RCM (2001) Syndromes of the head and neck. New York: Oxford University Press.

2. Heike CL, Hing AV (2009) Craniofacial Microsomia Overview. In: Pagon RA, Adam MP, Bird TD, Dolan CR, Fong CT et al., editors. GeneReviews. Seattle (WA).

3. Rimoin DL EA (2006) Emery and Rimoin’s principles and practice of medical genetics. Philadelphia: Churchill Livingstone.

4. Poswillo D (1988) The aetiology and pathogenesis of craniofacial deformity. Development 103 Suppl: 207-212.

5. Werler MM, Sheehan JE, Hayes C, Padwa BL, Mitchell AA, et al. (2004) Demographic and reproductive factors associated with hemifacial microsomia. Cleft Palate Craniofac J 41: 494–450.

6. Vendramini-Pittoli S, Kokitsu-Nakata NM (2009) Oculoauriculovertebral spectrum: report of nine familial cases with evidence of autosomal dominant inheritance and review of the literature. Clin Dysmorphol 18: 67–77.

7. Rollnick BR, Kaye CI (1983) Hemifacial microsomia and variants: pedigree data. Am J Med Genet 15: 233–253.

8. Cohen MM, Jr., Rollnick BR, Kaye CI (1989) Oculoauriculovertebral spectrum: an updated critique. Cleft Palate J 26: 276–286.

9. Kaye CI, Martin AO, Rollnick BR, Nagatoshi K, Israel J, et al. (1992) Oculoauriculovertebral anomaly: segregation analysis. Am J Med Genet 43: 913–917.

10. Dabir TA, Morrison PJ (2006) Trisomy 10p with clinical features of facio-auriculo-vertebral spectrum: a case report. Clin Dysmorphol 15: 25–27.

11. de Ravel TJ, Legius E, Brems H, Van Hoestenberghe R, Gillis PH, et al. (2001) Hemifacial microsomia in two patients further supporting chromosomal mosaicism as a causative factor. Clin Dysmorphol 10: 263–267.

12. Hodes ME, Gleiser S, DeRosa GP, Yune HY, Girod DA, et al. (1981) Trisomy 7 mosaicism and manifestations of Goldenhar syndrome with unilateral radial hypoplasia. J Craniofac Genet Dev Biol 1: 49–55.

13. Kobrynski L, Chitayat D, Zahed L, McGregor D, Rochon L, et al. (1993) Trisomy 22 and facioauriculovertebral (Goldenhar) sequence. Am J Med Genet 46: 68–71.

14. Miller R, Stephan MJ, Hume RF, Walker WO, Kelly P, et al. (2001) Extreme elevation of maternal serum alpha-fetoprotein associated with mosaic trisomy 8 in a liveborn. Fetal Diagn Ther 16: 120–122.

15. Pridjian G, Gill WL, Shapira E (1995) Goldenhar sequence and mosaic trisomy 22. Am J Med Genet 59: 411–413.

16. Kelberman D, Tyson J, Chandler DC, McInerney AM, Slee J, et al. (2001) Hemifacial microsomia: progress in understanding the genetic basis of a complex malformation syndrome. Hum Genet 109: 638–645.

17. Huang XS, Li X, Tan C, Xiao L, Jiang HO, et al. (2010) Genome-wide scanning reveals complex etiology of oculo-auriculo-vertebral spectrum. Tohoku J Exp Med 222: 311–318.

18. Rooryck C, Souakri N, Cailley D, Bouron J, Goizet C, et al. (2010) Array-CGH analysis of a cohort of 86 patients with oculoauriculovertebral spectrum. Am J Med Genet A 152A: 1984–1989.

19. Vermeesch JR, Balikova I, Schrander-Stumpel C, Fryns JP, Devriendt K (2011) The causality of de novo copy number variants is overestimated. Eur J Hum Genet 19: 1112–1113.

20. Aizenbud D, Shoham NV, Constantini S, Nevo N, Ben Arush M, et al. (2013) Goldenhar syndrome and medulloblastoma: A coincidental association? The first case report. J Craniomaxillofac Surg.

21. Kiezun A, Garimella K, Do R, Stitziel NO, Neale BM, et al. (2012) Exome sequencing and the genetic basis of complex traits. Nat Genet 44: 623–630.

22. Marth GT, Yu F, Indap AR, Garimella K, Gravel S, et al. (2011) The functional spectrum of low-frequency coding variation. Genome Biol 12: R84.

23. Ng SB, Turner EH, Robertson PD, Flygare SD, Bigham AW, et al. (2009) Targeted capture and massively parallel sequencing of 12 human exomes. Nature 461: 272–276.

24. (2002) The NCBI Handbook. In: McEntyre J, Ostell J, editors. Bethesda, MD: National Center for Biotechnology Information.

25. Hashimoto A, Hashimoto S, Ando R, Noda K, Ogawa E, et al. (2011) GEP100-Arf6-AMAP1-cortactin pathway frequently used in cancer invasion is activated by VEGFR2 to promote angiogenesis. PLoS One 6: e23359.

26. Akiyama K, Narita A, Nakaoka H, Cui T, Takahashi T, et al. (2010) Genome-wide association study to identify genetic variants present in Japanese patients harboring intracranial aneurysms. J Hum Genet 55: 656–661.

27. Maglott D, Ostell J, Pruitt KD, Tatusova T (2011) Entrez Gene: gene-centered information at NCBI. Nucleic Acids Res 39: D52–57.

28. Itsara A, Cooper GM, Baker C, Girirajan S, Li J, et al. (2009) Population analysis of large copy number variants and hotspots of human genetic disease. Am J Hum Genet 84: 148–161.

29. Bray SM, Mulle JG, Dodd AF, Pulver AE, Wooding S, et al. (2010) Signatures of founder effects, admixture, and selection in the Ashkenazi Jewish population. Proc Natl Acad Sci U S A 107: 16222–16227.

30. Genomes Project C, Abecasis GR, Auton A, Brooks LD, DePristo MA, et al. (2012) An integrated map of genetic variation from 1,092 human genomes. Nature 491: 56–65.

31. Ou Z, Martin DM, Bedoyan JK, Cooper ML, Chinault AC, et al. (2008) Branchiootorenal syndrome and oculoauriculovertebral spectrum features associated with duplication of SIX1, SIX6, and OTX2 resulting from a complex chromosomal rearrangement. Am J Med Genet A 146A: 2480–2489.

32. Northup JK, Matalon D, Hawkins JC, Matalon R, Velagaleti GV (2010) Pericentric inversion, inv(14)(p11.2q22.3), in a 9-month old with features of Goldenhar syndrome. Clin Dysmorphol 19: 185–189.

33. Ballesta-Martinez MJ, Lopez-Gonzalez V, Dulcet LA, Rodriguez-Santiago B, Garcia-Minaur S, et al. (2013) Autosomal dominant oculoauriculovertebral spectrum and 14q23.1 microduplication. Am J Med Genet A 161: 2030–2035.

34. Moreau Y, Tranchevent LC (2012) Computational tools for prioritizing candidate genes: boosting disease gene discovery. Nat Rev Genet 13: 523–536.

35. Erlich Y, Edvardson S, Hodges E, Zenvirt S, Thekkat P, et al. (2011) Exome sequencing and disease-network analysis of a single family implicate a mutation in KIF1A in hereditary spastic paraparesis. Genome Res 21: 658–664.

36. Oti M, Brunner HG (2007) The modular nature of genetic diseases. Clin Genet 71: 1–11.

37. van Driel MA, Bruggeman J, Vriend G, Brunner HG, Leunissen JA (2006) A text-mining analysis of the human phenome. Eur J Hum Genet 14: 535–542.

38. Keegan CE, Mulliken JB, Wu BL, Korf BR (2001) Townes-Brocks syndrome versus expanded spectrum hemifacial microsomia: review of eight patients and further evidence of a “hot spot” for mutation in the SALL1 gene. Genet Med 3: 310–313.

39. Kallen K, Robert E, Castilla EE, Mastroiacovo P, Kallen B (2004) Relation between oculo-auriculo-vertebral (OAV) dysplasia and three other non-random associations of malformations (VATER, CHARGE, and OEIS). Am J Med Genet A 127A: 26–34.

40. Aerts S, Lambrechts D, Maity S, Van Loo P, Coessens B, et al. (2006) Gene prioritization through genomic data fusion. Nat Biotechnol 24: 537–544.

41. Chen J, Bardes EE, Aronow BJ, Jegga AG (2009) ToppGene Suite for gene list enrichment analysis and candidate gene prioritization. Nucleic Acids Res 37: W305–311.

42. Maurano MT, Humbert R, Rynes E, Thurman RE, Haugen E, et al. (2012) Systematic localization of common disease-associated variation in regulatory DNA. Science 337: 1190–1195.

43. Lage K, Hansen NT, Karlberg EO, Eklund AC, Roque FS, et al. (2008) A large-scale analysis of tissue-specific pathology and gene expression of human disease genes and complexes. Proc Natl Acad Sci U S A 105: 20870–20875.

44. Huang N, Lee I, Marcotte EM, Hurles ME (2010) Characterising and predicting haploinsufficiency in the human genome. PLoS Genet 6: e1001154.

45. Matsuo I, Kuratani S, Kimura C, Takeda N, Aizawa S (1995) Mouse Otx2 functions in the formation and patterning of rostral head. Genes Dev 9: 2646–2658.

46. Hide T, Hatakeyama J, Kimura-Yoshida C, Tian E, Takeda N, et al. (2002) Genetic modifiers of otocephalic phenotypes in Otx2 heterozygous mutant mice. Development 129: 4347–4357.

47. Fossat N, Chatelain G, Brun G, Lamonerie T (2006) Temporal and spatial delineation of mouse Otx2 functions by conditional self-knockout. EMBO Rep 7: 824–830.

48. Chassaing N, Sorrentino S, Davis EE, Martin-Coignard D, Iacovelli A, et al. (2012) OTX2 mutations contribute to the otocephaly-dysgnathia complex. J Med Genet 49: 373–379.

49. Nolen LD, Amor D, Haywood A, St Heaps L, Willcock C, et al. (2006) Deletion at 14q22-23 indicates a contiguous gene syndrome comprising anophthalmia, pituitary hypoplasia, and ear anomalies. Am J Med Genet A 140: 1711–1718.

50. Diaczok D, Romero C, Zunich J, Marshall I, Radovick S (2008) A novel dominant negative mutation of OTX2 associated with combined pituitary hormone deficiency. J Clin Endocrinol Metab 93: 4351–4359.

51. Bunt J, Hasselt NE, Zwijnenburg DA, Hamdi M, Koster J, et al. (2012) OTX2 directly activates cell cycle genes and inhibits differentiation in medulloblastoma cells. Int J Cancer 131: E21–32.

52. Northcott PA, Korshunov A, Witt H, Hielscher T, Eberhart CG, et al. (2011) Medulloblastoma comprises four distinct molecular variants. J Clin Oncol 29: 1408–1414.

53. Blumenstiel B, Cibulskis K, Fisher S, DeFelice M, Barry A, et al. (2010) Targeted exon sequencing by in-solution hybrid selection. Curr Protoc Hum Genet Chapter 18: Unit 18 14.

54. Pruitt KD, Harrow J, Harte RA, Wallin C, Diekhans M, et al. (2009) The consensus coding sequence (CCDS) project: Identifying a common protein-coding gene set for the human and mouse genomes. Genome Res 19: 1316–1323.

55. Langmead B, Trapnell C, Pop M, Salzberg SL (2009) Ultrafast and memory-efficient alignment of short DNA sequences to the human genome. Genome Biol 10: R25.

56. Li H, Durbin R (2009) Fast and accurate short read alignment with Burrows-Wheeler transform. Bioinformatics 25: 1754–1760.

57. Li H, Handsaker B, Wysoker A, Fennell T, Ruan J, et al. (2009) The Sequence Alignment/Map format and SAMtools. Bioinformatics 25: 2078–2079.

58. Koboldt DC, Chen K, Wylie T, Larson DE, McLellan MD, et al. (2009) VarScan: variant detection in massively parallel sequencing of individual and pooled samples. Bioinformatics 25: 2283–2285.

59. McKenna A, Hanna M, Banks E, Sivachenko A, Cibulskis K, et al. (2010) The Genome Analysis Toolkit: a MapReduce framework for analyzing next-generation DNA sequencing data. Genome Res 20: 1297–1303.

60. Gymrek M, Golan D, Rosset S, Erlich Y (2012) lobSTR: A short tandem repeat profiler for personal genomes. Genome Res 22: 1154–1162.

61. Rozen S, Skaletsky H (2000) Primer3 on the WWW for general users and for biologist programmers. Methods Mol Biol 132: 365–386.

62. Fujita PA, Rhead B, Zweig AS, Hinrichs AS, Karolchik D, et al. (2011) The UCSC Genome Browser database: update 2011. Nucleic Acids Res 39: D876–882.

63. Korn JM, Kuruvilla FG, McCarroll SA, Wysoker A, Nemesh J, et al. (2008) Integrated genotype calling and association analysis of SNPs, common copy number polymorphisms and rare CNVs. Nat Genet 40: 1253–1260.

64. Saccone SF, Quan J, Jones PL (2012) BioQ: tracing experimental origins in public genomic databases using a novel data provenance model. Bioinformatics 28: 1189–1191.

65. Quinlan AR, Hall IM (2010) BEDTools: a flexible suite of utilities for comparing genomic features. Bioinformatics 26: 841–842.

66. Purcell S, Neale B, Todd-Brown K, Thomas L, Ferreira MA, et al. (2007) PLINK: a tool set for whole-genome association and population-based linkage analyses. Am J Hum Genet 81: 559–575.

67. Abecasis GR, Cherny SS, Cookson WO, Cardon LR (2002) Merlin--rapid analysis of dense genetic maps using sparse gene flow trees. Nat Genet 30: 97–101.

68. Barrett T, Wilhite SE, Ledoux P, Evangelista C, Kim IF, et al. (2013) NCBI GEO: archive for functional genomics data sets--update. Nucleic Acids Res 41: D991–995.

